# Reversibility of resistance in a fluctuation test experiment modifies the tail of the Luria-Delbrück distribution

**DOI:** 10.1101/2022.12.26.521941

**Authors:** Pavol Bokes, Anna Hlubinová, Abhyudai Singh

**Affiliations:** Department of Applied Mathematics and Statistics, Comenius University, 84248 Bratislava, Slovakia; Departments of Electrical and Computer Engineering, Biomedical Engineering, University of Delaware, Newark, DE 19716

**Keywords:** Luria–Delbrück fluctuation test, reversible state switching, back mutation, mutation rate, Landau distribution, mathematical biology

## Abstract

We consider a fluctuation test experiment in which cell colonies are grown from a single cell until they reach a given population size, and then they are exposed to treatment. While they grow, the cells may, with a low probability, acquire resistance to treatment and pass it on to their offspring. Unlike the classical Luria–Delbrück fluctuation test and motivated by recent work on drug-resistance acquisition in cancer/microbial cells, we allow for the resistant cell state to switch back to a drug-sensitive state. This modification does not affect the central part of the (Luria–Delbrück) distribution of the number of resistant survivors: the previously developed approximation by the Landau probability density function applies. However, the right tail of the modified distribution deviates from the power law decay of the Landau distribution. We demonstrate that the correction factor is equal to the Landau cumulative distribution function.

## 1. Introduction

In a fluctuation experiment, a colony of bacterial/cancer cells is grown, and eventually exposed to a treatment [1]. Most cells are killed, but some are resistant to the treatment. Repeated experiments each give a different number of resistant cells. The variability in the number of survivors is key in a fluctuation test. If the resistance is acquired when the treatment is administered, a low Poissonian variability is expected; on the other hand, if it can be acquired by a cell at an early stage of the colony growth, and passed on to its (many) descendants, the variability will be large. The resistant cells are typically assumed to have developed a genetic mutation, and are referred to as “mutants”. However, several studies have shown that stochastic expression of specific proteins can lead to resistance to drug therapy arising *transiently*. In such case, cells switch to become resistant for several generations, but then switch back into a drug-sensitive state [2–6]. Several tools that exploit the colony-to-colony variation to estimate the rates of switching into and out of the resistant state have been recently developed [7,8].

Mathematical analysis has been applied, now for decades, to quantify this variability under different modelling assumptions and parameter regimes [9]. The modelling framework is either semi-deterministic or fully stochastic [10]. In the former case, proliferation is deterministic and the acquisition of resistance is stochastic [11], [12]; in the latter case both are stochastic. The fully stochastic framework further differentiates into fixed-time or fixed-population statistical ensembles. In the former case, one waits until a given end time and then takes the measurement of the cell population [13]. The final measurement will include the effect of total population noise. In the fixed-population case, the measurement is taken when the total cell population reaches a given level [14], [15]. Such an approach excludes the population noise, and is thus consistent with the semi-deterministic framework. An alternative approach is to look at the proportion of resistant cells rather than the absolute number [8].

In most models, resistance, once acquired, is never lost: resistant mothers bear resistant daughters. Such models are asymmetric, whereas ones in which bidirectional change is possible are structurally symmetric. The possibility of a “back mutation” is suggested in [16] and modelled deterministically in [17]. Back-mutation is inherently present in a recent innovative model which combines population growth with mutations in a sequence of nucleotides [18]. The loss of resistance by random changes in gene expression has been suggested to reduce variability in a fluctuation experiment [2,7,8].

The aim of this paper is to understand the probability distribution of the number of resistant cells in a structurally symmetric, fully-stochastic, fixed-population-ensemble fluctuation test. Some explicit results are available in the asymmetric case but they reveal little [14]. In the asymptotic regime of small probability of resistance acquisition, the Luria–Delbrück distribution of the number of surviving resistors can be approximated by the (discrete) Lea–Coulson distribution [19], [14], [10]. The Lea–Coulson distribution has a shape parameter that corresponds to the population rate of resistance acquisition. If this is sufficiently large, the (discrete) Lea–Coulson distribution simplifies to its coarse-grained continuous counterpart, the Landau distribution [20], [21], [10]. The Landau distribution is a one-sided stable distribution with characteristic exponent one [22]. The Landau approximation is universal (or self-similar) because the resistance rate only affects its position and scale parameters. Both Lea–Coulson and Landau distributions have a heavy tail. Our paper contributes by characterising the correction to the power-law tail that is due to loss of resistance.

The Lea–Coulson/Landau approximations represent, in the parlance of matched asymptotics, a boundary-layer approximation to the Luria–Delbrück distribution. They are valid provided that the overwhelming majority of cells are sensitive. The boundary layer solution can be combined with a regular power series solution to construct a composite solution that approximates the Luria–Delbrück distribution uniformly [10]. The reversal of resistance introduces a new boundary layer solution that is valid if the overwhelming majority of cells are resistant. The uniform solution will combine the regular power series solution as well as the two boundary layer solutions. Our analysis will thus extend the matched asymptotic framework of [10].

## 2. Model formulation

The population, which is structured into sensitive and resistant cells, proliferates as per a Markov jump chain. Each step grows the population by one. Whether the new cell is sensitive or resistant is determined probabilistically depending on a randomly chosen mother cell (Fig 1). Thus, after each step, the number of resistant cells either stays the same or increases by one; the probabilities of these two alternative transitions are given in Fig 2a. The total population size *N* is thereby used as the independent variable; the number of resistant cells *m* = *m*(*N*) is a time-inhomogeneous Markov chain in discrete ‘time’ *N*.

**Figure 1.**
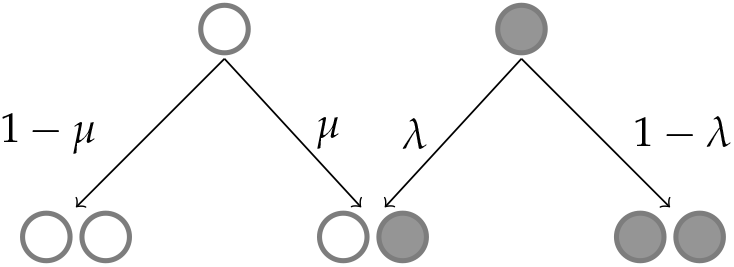
White balls in the diagram represent sensitive cells, while black balls represent resistant cells. Arrows pointing from top tier balls (mother cells) downwards illustrate the four possible outcomes of cell division: left, a sensitive parent cell may either have a sensitive descendant or its offspring may be resistant, with probabilities 1 − *μ* and *μ*, respectively; right, a resistant cell may also produce a sensitive or resistant offspring, with probabilities *λ* and 1 − *λ*, respectively.

**Figure 2.**
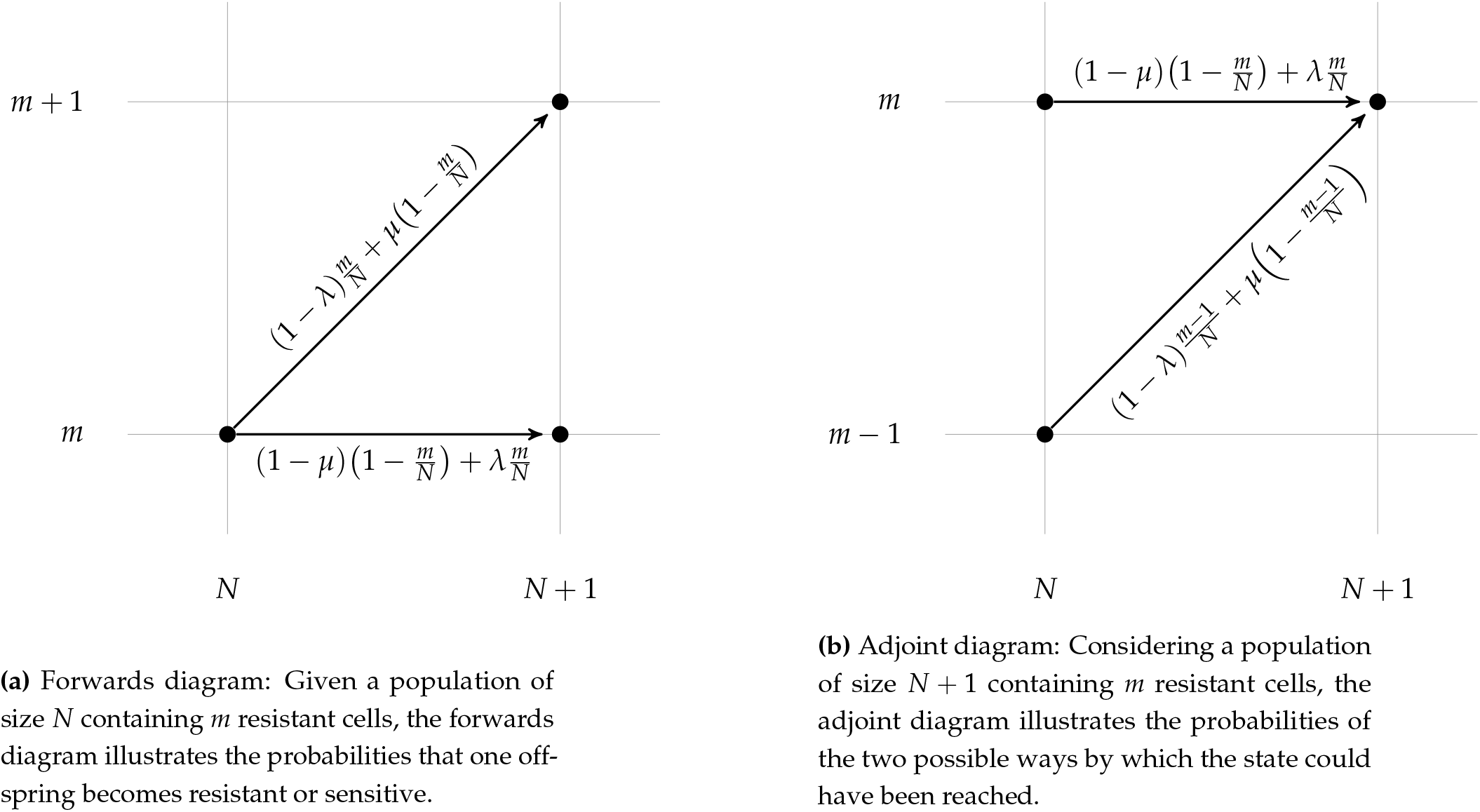
Forwards and adjoint schematic illustrations of transition probabilities in the Markov chain model. The parameters *μ* and *λ* are the resistance acquisition and loss probabilities, respectively.

The probability *P*_*N*_(*m*) of having *m* resistant cells in a total population of *N* cells satisfies the master equation

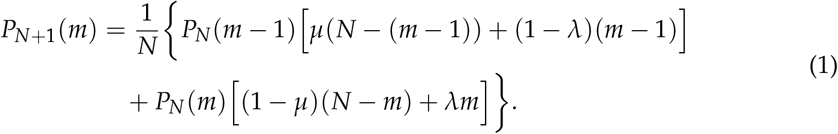

The bracketed terms in (1) are the transition probabilities of the adjoint diagram of the process (Fig 2b). For *λ* = 0, (1) simplifies to the master equation for the irreversible case as previously studied in [10]. Further details can be found in Appendix A.

The master equation (1) is studied subject to boundary conditions

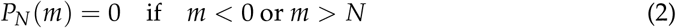

and an initial condition

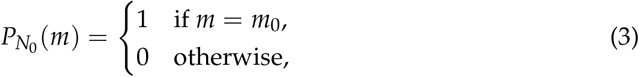

wherein *N*_0_ stands for the initial population size and *m*_0_ for the initial resistant subpopulation size.

## 3. Methods

Following previous studies [10,11], as well as empirical evidence [2], we assume that the resistance acquisition and loss probabilities are small perturbation parameters, *μ* ≪ 1 and *λ* ≪ 1. The aim of this paper is to characterise the asymptotic behaviour of the solution to the master equation (1) subject to (2) and (3) as the perturbation parameters tend to zero. We thereby assume that they tend to zero with the same rate: *μ*/*λ* = *O*(1).

It turns out that the problem in question is singularly perturbed and there are several alternative distinguished approximations that apply in different regions of the definition domain 0 ≤ *m* ≤ *N* < ∞ of the solution *P*_*N*_(*m*), as well as intermediate approximations through which the former can be matched. Additionally, the analysis proceeds differently depending on whether we start with a homogeneous population, i.e. with *m*_0_ = 0 resistant and *N*_0_ > 0 sensitive cells or with *m*_0_ = *N*_0_ > 0 resistant cells, or we start with a mixed population with *m*_0_ > 0 resistant and *N*_0_ − *m*_0_ > 0 sensitive cells. Our aim is to treat all the possible combinations exhaustively; to help the reader navigate through the resulting complexity, Table 1 outlines the region of validity for the individual approximations and gives references to their formulae. Below we provide a brief commentary on the table. The derivations are found in subsections that follow.

**Table 1.**
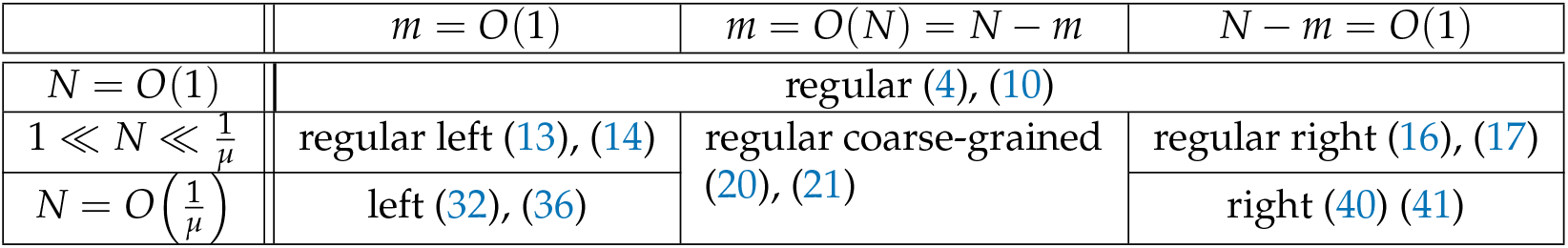
Overview of approximate solutions to the equation (1) and their regions of validity within the definition domain 0 ≤ *m* ≤ *N* < ∞. The first reference points to a formula that applies for a mixed initial population; the second reference points to an exclusively sensitive initial population. The linked formulae contain the population transition rates (23) and the (partial summations of the) Lea–Coulson probability mass function (33)–(34).

| | $m = O(1)$ | $m = O(N) = N - m$ | $N - m = O(1)$ |
| --- | --- | --- | --- |
| $N = O(1)$ | regular (4), (10) | | |
| $1 \ll N \ll \frac{1}{\mu}$ | regular left (13), (14) | regular coarse-grained<br>(20), (21) | regular right (16), (17) |
| $N = O\left(\frac{1}{\mu}\right)$ | left (32), (36) | | right (40) (41) |

The first asymptotic approximation is developed under the assumption of bounded population sizes *N* = *O*(1) as the perturbation parameters tend to zero. The number of resistant cells is then also bounded via the natural constraint *m* ≤ *N*. Mathematically, the solution is developed into a regular power series in the perturbation parameter and approximated by the leading order term. Therefore, we call it a regular approximation to the solution or simply regular solution. The approximation occupies the first row of Table 1 and is derived in Section 3.1.

The regular approximation is an example of a distinguished limit: the variables are of a definite order *O*(1). Another distinguished limit occurs if the total population size is inversely proportional to the small perturbation parameter and the number of resistant cells is *O*(1) (Table 1, row 3, column 1). This is the large population and low mutation rate scenario discussed in previous works on the unidirectional case [10,11]. Our structurally symmetric can also develop a right boundary layer occurring at *O*(1) numbers of sensitive cells (Table 1, row 3, column 3).

Three intermediate limits are obtained by taking the total population large in the regular solution and considering various scales of the resistant and sensitive subpopulations (Table 1, row 2; Sections 3.2–3.4). The intermediate left and right regular solutions are used for the asymptotic matching of the distinguished left and right solutions (Sections 3.5 and 3.6). The coarse-grained regular solution is applicable in the space between the two boundary layers (Table 1, row 3, column 2) and can be combined with the layer solutions into a uniform composite approximation (Section 3.7).

The triangular character of the definition domain in Table 1 renders the concept of inner and outer solutions ambiguous. On the one hand, following [10], one can refer to the boundary layer solutions as inner because they focus on small fractions of the total population. On the other hand, following classical applications of singular perturbations in biology [23], one can also refer to the regular solution as inner because it captures the system’s early dynamics. In order to avoid a clash with earlier conventions, we decide not to use the names inner and outer solutions in this work.

### 3.1. Regular solution

By a regular solution to the master equation (1) we understand a solution that depends on the perturbation parameter *μ* regularly, i.e. is a power series in *μ*. Our aim is to determine this solution at the leading order, i.e. provide the first non-zero term of the power-series. The case *m*_0_ > 0 and *N*_0_− *m*_0_ > 0 (both resistant and sensitive cells present initially) is, perhaps surprisingly, easier than the case *m*_0_ = 0, *N*_0_ > 0 or the symmetrically opposite case of *m*_0_ = *N*_0_ > 0, and will be treated first.

#### 3.1.1. Case *m*_0_ > 0 and *N*_0_ − *m*_0_ > 0

Rather than formally solving the difference equation (1), it is both easier and more instructive to derive the regular solution from probabilistic considerations. The leading-order behaviour of the regular solution is obtained simply by neglecting the perturbations *μ* and *λ* in (1). The resulting equation describes a growing culture of cells which deterministically pass their state to their progeny. The growth goes as follows: one picks a cell at random and looks at its state; the chosen cell is returned to the culture as well as another cell (a daughter) is added that has the same state.

Aside from the trivia of terminology (cells = balls; state = colour), this is a classical Polya urn model [24]. Detailed description of this model can be found in Appendix B. As a result, with *μ* (and *λ*) approaching zero,

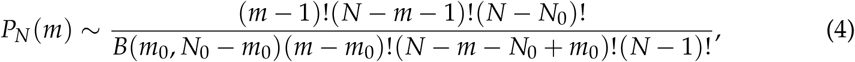

where *m*_0_ ≤ *m* ≤ *N*− *N*_0_ and the tilde sign is understood in the usual sense of asymptotic expansions [25]. *B*(*x, y*) = Γ(*x*)Γ(*y*)/Γ(*x* + *y*) = (*x*− 1)!(*y*− 1)!/(*x* + *y* −1)! is the beta function.

#### 3.1.2. Case *m*_0_ = 0, *N*_0_ > 0

The case of an entirely sensitive initial population is of the prime interest in the paper. It is more difficult than the mixed initial population case in Section 3.1.1. We will make use of the previous case in deriving the new result. First, note that neglecting the perturbations *μ* and *λ* in (1) leads to a problem with a trivial solution. Namely, one always picks a sensitive cell and keeps adding new sensitive cells. Mathematically,

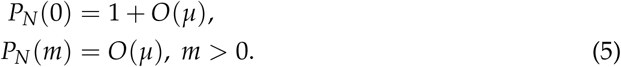

The second equation in (5) tells us that probability of finding some sensitive cells is small, but does not describe the leading-order — here O(*μ*) — behaviour. For this, we need a subtler estimate, which we will obtain, once again, from probabilistic considerations rather than formal solution techniques for difference equations.

Let *L* denote the population size such that the first *L* cells are sensitive and the (*L* + 1) - th cell is resistant. Conditioning on the value of *L* we have

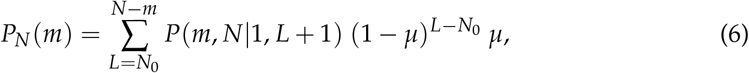

where *P*(*m, N* | 1, *L* + 1) denotes the probability of observing *m* resistant cells in a population of size *N* that was initially of size *L* + 1 and contained one resistant cell. This can be estimated by setting *m*_0_ = 1 *N*_0_ = *L* + 1 into (4), which yields

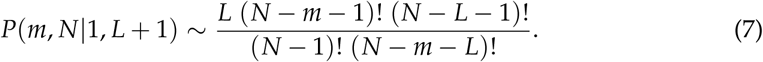

Since the main aim is to derive a leading-order approximation of *P*_*N*_(*m*), one may neglect *μ* in the factors (1 − *μ*) appearing in (6). As a result we get

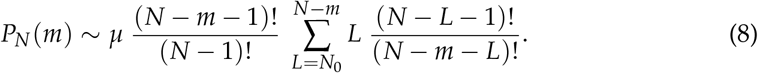

Moreover, using symbolic software (details in Appendix C), the sum appearing in the previous equation can be simplified

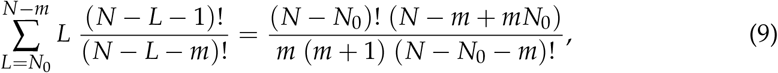

giving final result

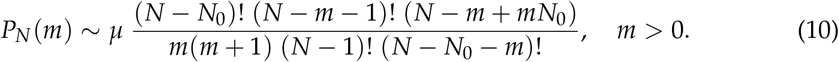

Formula (10) was derived by other means for the unidirectional model in [10].

#### 3.1.3. Case *m*_0_ = *N*_0_ > 0

This case is a mirror reflection of the case discussed in Section 3.1.2. The result, which reads *P*_*N*_(*N*) ∼ 1 and

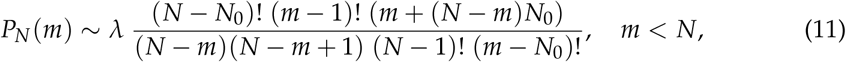

is readily obtained by replacing *μ* by *λ, N*_0_ − *m*_0_ by *m*_0_, *N* − *m* by *m*, and vice versa, in (10).

### 3.2. Regular left solution

By a regular left solution we mean the approximate solution that considers relatively large total population, 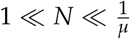, which includes only a small number of resistant cells *m* = *O*(1). The regular left solution can be derived from the regular solution formulae (4), (10), and (11) by approximating descending factorials by powers as appropriate; for example, one has

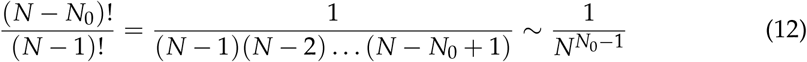

for *N* ≫ 1.

#### 3.2.1. Case *m*_0_ > 0 and *N*_0_ − *m*_0_ > 0

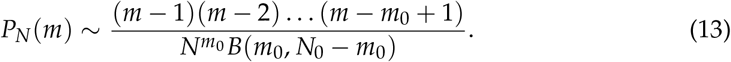

#### 3.2.2. Case *m*_0_ = 0, *N*_0_ > 0

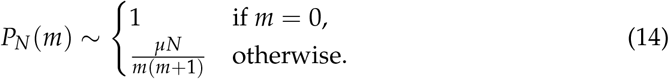

#### 3.2.3. Case *m*_0_ = *N*_0_ > 0

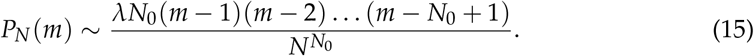

### 3.3. Regular right solution

By a regular right solution we mean the approximate solution that considers relatively large total population, 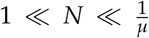, which includes a large number of resistant cells, mathematically *N* − *m* = *O*(1). These approximations are most conveniently obtained by performing substitutions *m* ↔ *N* − *m, m*_0_ ↔ *N*_0_ − *m*_0_, and *λ* ↔ *μ* in (13), (14), and (15).

#### 3.3.1. Case *m*_0_ > 0 and *N*_0_ − *m*_0_ > 0

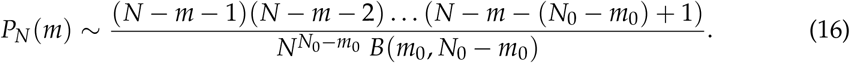

#### 3.3.2. Case *m*_0_ = 0, *N*_0_ > 0

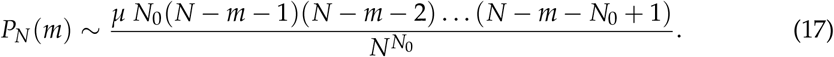

#### 3.3.3. Case *m*_0_ = *N*_0_ > 0

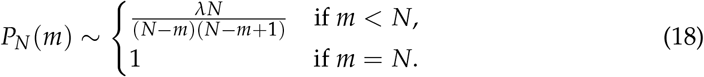

### 3.4. Regular coarse-grained solution

By a regular coarse-grained solution we mean an approximate solution that considers a large number of cells in a total population, while the order of the number of resistant and sensitive cells in the population is the same. Mathematically *m* = *O*(*N*) = *N* − *m*. The regular coarse-grained solution is obtained, similarly as regular left/right solutions, by approximating descending factorials by appropriate powers in the regular solution formulae (4), (10), and (11). For example,

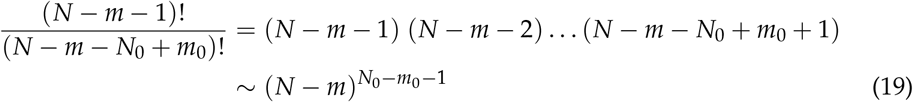

holds for *m* and *N* both large and of the same order.

#### 3.4.1. Case *m*_0_ > 0 and *N*_0_ − *m*_0_ > 0

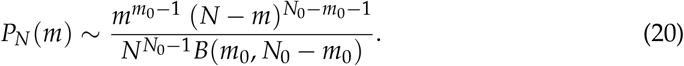

#### 3.4.2. Case *m*_0_ = 0, *N*_0_ > 0

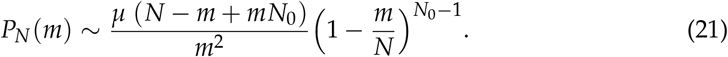

#### 3.4.3. Case *m*_0_ = *N*_0_ > 0

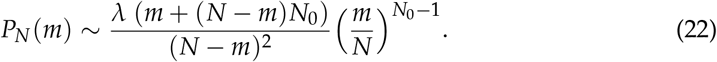

We note that (20) relates to the classical convergence result to the beta distribution for the Polya urn [26] and that (21) and (22) are symmetric.

### 3.5. Left boundary layer solution

As in the previous sections, we assume that the resistance acquisition and loss probabilities are small, i.e. *μ* ≪ 1 and *λ* ≪ 1 such that *λ*/*μ* = *O*(1), and that the total population size is large, i.e. *N* ≫ 1. In contrast with the previous sections, here we focus on the distinguished large-population low-mutation-rate regime in which the population size is as large as to make the population rates of resistance acquisition and loss,

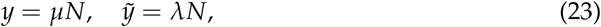

of order one. The distinguished limit will be found for *m* = *O*(1) resistant cells. This represents a (left) boundary layer of the large interval of admissible values *m ∈* {0, 1, …, *N*}. The right boundary layer, occurring for *N*− *m* = *O*(1), will be treated in the next section by symmetry arguments. Unless stated otherwise, the asymptotic equivalence sign ∼ will used in the section the context of the limit process *μ* → 0, *y* = *O*(1), 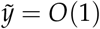, *m* = *O*(1).

The boundary-layer solution is found by the method of the generating function, which is defined via

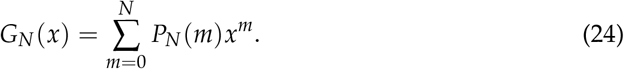

The usual rules [27] transform the master equation (1) into a difference–differential equation

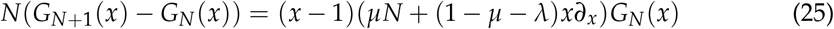

for the generating function.

Guided by the intermediate asymptotics (13), (14), and (15) of the regular solution, we look for a generating function with the same kind of asymptotics,

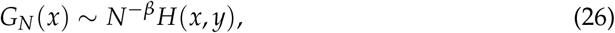

where *β* is to be matched to regular solution. Note that

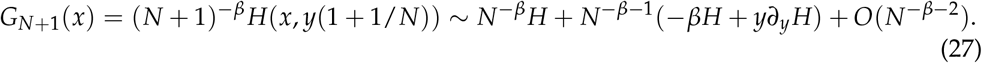

Inserting (26) and (27) into (25) and collecting leading order terms, we obtain the partial differential equation

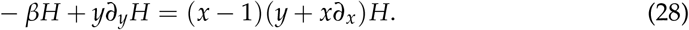

For *β* = 0, (28) is known as the Lea–Coulson equation [9]. The parameter *β* can be viewed as an eigenvalue and the solution *H* as an eigenfunction of the Lea–Coulson differential operator.

Solving (28) by the method of characteristics, one uncovers a parametric family of solutions

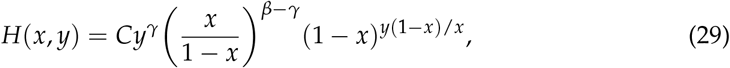

where *C* and *γ* are arbitrary constants. In what follows, we determine these constants and the eigenvalue *β* by matching to the regular solution. The distinct cases introduced previously are thereby treated separately.

#### 3.5.1. Case *m*_0_ > 0, *N*_0_ − *m*_0_ > 0

The left–regular approximation (13), when expressed in terms of the generating function, reads

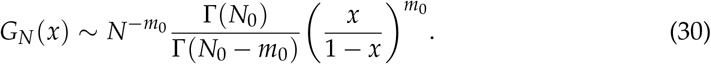

The approximation is valid in the left boundary layer *m* = *O*(1) for an intermediate range of population sizes 1 ≪ *N* ≪ 1/*μ*.

Matching (30) with (26) and (29) yields *γ* = 0, *β* = *m*_0_, and *C* = Γ(*N*_0_)/Γ(*N*_0_ − *m*_0_), i.e.

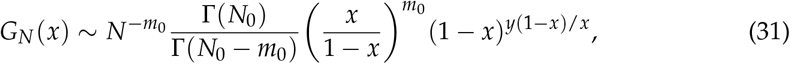

which is valid in the left boundary layer *m* = *O*(1) for population sizes extending up to *N* = *O*(1/*μ*).

The right-hand side of (31) can be transformed into the probability space by combining the following rules

- *x* transforms to the shift operator;
- (1 −*x*)^−1^ transforms to the summation operator (a convolution with a sequence of ones);
- (1 − *x*)^*y*(1−*x*)/*x*^ transforms to the Lea-Coulson probability mass function (pmf).

The approximation (31), when expressed in terms of the probability mass function, therefore reads

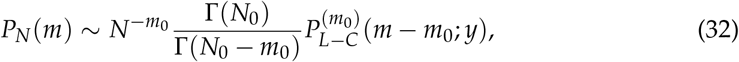

where 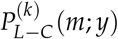 is the Lea–Coulson probability mass function *k* times summed over, i.e.

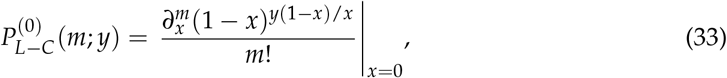

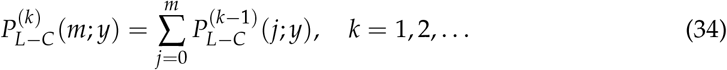

In particular, 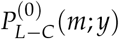 is the Lea–Coulson pmf and 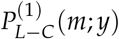 is the Lea–Coulson cumulative distribution function (cdf).

#### 3.5.2. Case *m*_0_ = 0, *N*_0_ > 0

The left-regular intermediate approximation (14) implies *G*_*N*_(*x*) ∼1 for 1 ≪ *N* ≪ 1/*μ*, which when matched to (29) and (26) implies *β* = *γ* = 0, *C* = 1; the left boundary approximation then reads

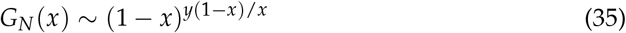

in the generating function space and

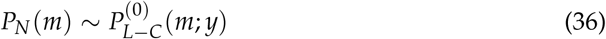

in the probability space. In this case, the matching procedure is equivalent to directly imposing the initial condition on the solution to the Lea–Coulson equation [10].

#### 3.5.3. Case *m*_0_ = *N*_0_ > 0

The intermediate approximation (15) implies

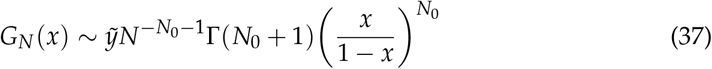

for 1 ≪ *N* ≪ 1/*μ*. Matching with (29) and (26) yields *γ* = 1, *β* = *N*_0_ + 1, *C* = Γ(*N*_0_ + *λ*/*μ*, i.e.

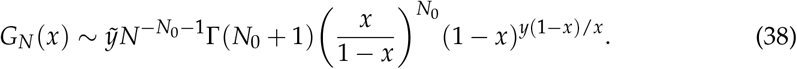

This reads

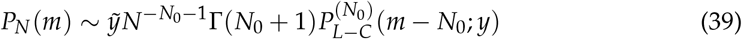

in the probability space.

### 3.6. Right boundary layer solution

Symmetry considerations imply that 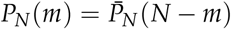, where the bar indicates a model in which the substitutions *m*_0_ *N*_0_ − *m*_0_ and 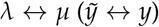 are made. Combining the symmetry result with the previous section’s results on the left boundary layer yield the following list of approximations. The equivalence sign is used in the context of *μ* → 0, *y* = *O*(1), 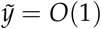, *N* − *m* = *O*(1).

#### 3.6.1. Case *m*_0_ > 0, *N*_0_ − *m*_0_ > 0

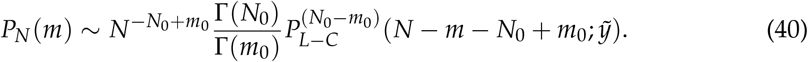

#### 3.6.2. Case *m*_0_ = 0, *N*_0_ > 0

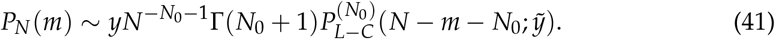

#### 3.6.3. Case *m*_0_ = *N*_0_ > 0

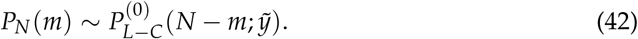

### 3.7. Log-composite solution

Composite solutions provide a means of combining multiple scale asymptotic approximations into a single uniformly valid approximation. In the most common example of a composite solution, the inner and outer solutions are added up and their common limit is subtracted [23]. In the log-composite solution, addition and subtraction is replaced by multiplication and division.

For *N* = *O*(1/*μ*), three approximations are on offer: the two boundary approximations and the (coarse-grained) regular approximation. Each of them is valid only on its own portion of the definition domain *m ∈*{ 0, 1, …, *N*} : the left solution applies for *m* = *O*(1), the regular coarse-grained for *m* = *O*(*N*) = *N* − *m*, and the right solution for *N*− *m* = *O*(1). Here we combine the individual asymptotic approximations to construct a uniformly valid composite approximation. The construction follows the pattern of

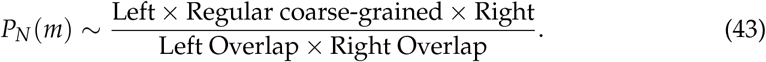

By “left overlap” and “right overlap” we mean the coarse-grained versions of the left and right regular solutions (Sections 3.2 and 3.3). If the variable *m* is in the range of validity of the left boundary solution, then the left overlap cancels with outer (and right overlap with the right solution). Similarly, if the variable *m* is in the range of the validity of the outer solution, the left overlap cancels with the left solution. The result is a uniform approximation across the entire interval of admissible values of *m*.

Application of the above principle in the usual cases leads to:

#### 3.7.1. Case *m*_0_ > 0, *N*_0_ − *m*_0_ > 0

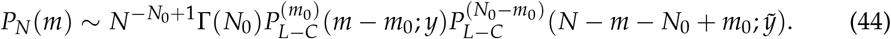

#### 3.7.2. Case *m*_0_ = 0, *N*_0_ > 0

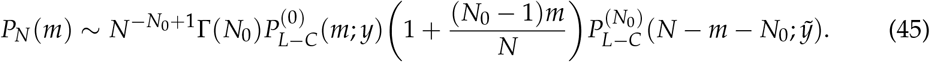

#### 3.7.3. Case *m*_0_ = *N*_0_ > 0

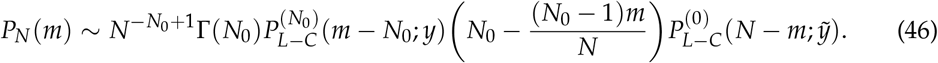

### 3.8. Landau distribution

As the shape parameter *y* of the Lea–Coulson pmf 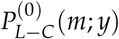 increases, the pmf can be approximated by the probability density function (pdf) of the Landau distribution [10]. Additionally, the partial summation 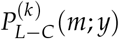, *k* = 1, 2, …, can be approximated by a partial integration of the pdf. Precisely,

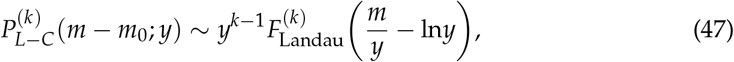

where

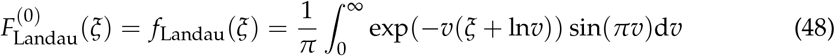

is the density of the Landau distribution [22,28] and

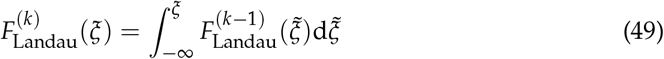

is the *k* th partial integration of the density. In particular, 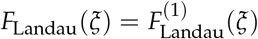is the cdf of the Landau distribution.

## 4. Results

In this section we collate the asymptotic analyses of Section 3 and extract the most significant results. We specifically focus on two relatively simple but important cases of initial condition in which the Landau cumulative distribution function appears: first, starting with a single sensitive cell and second, starting with two cells, one sensitive and one resistant. We then also discuss more complex deterministic as well as stochastic initial conditions. In numerical examples, we adopt the parameter values from the study of the unidirectional model [10]: the final total population is *N* = 2 *×* 10^5^ and the probability of resistance acquisition is *μ* = 10^−3^. We compare the reversible case, where we take the probability of resistance loss to be equal to that of resistance acquisition (*λ* = *μ*), with the previously studied unidirectional situation (*λ* = 0).

### 4.1. One sensitive cell at the beginning

This type of initial condition has been widely considered in studies concerning the Luria–Delbrück fluctuation test. Approximations of the resistant population distribution by the Lea–Coulson probability mass function and the Landau probability density functions have previously been reported. This paper establishes the appearance of partial summations/integrations of these distributions, in particular the cumulative distribution function, in the tail of the population distribution.

For low to moderate values *m*, the probability of seeing *m* resistant cells among *N* total cells is approximated by (see Figure 3a)

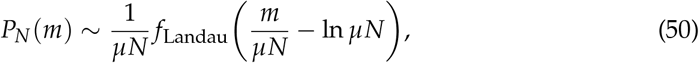

where *f*_Landau_(*ξ*) is the density (48) of the Landau distribution. The approximation follows from (36) and (47). This approximation was previously derived for the unidirectional model in [10].

**Figure 3.**
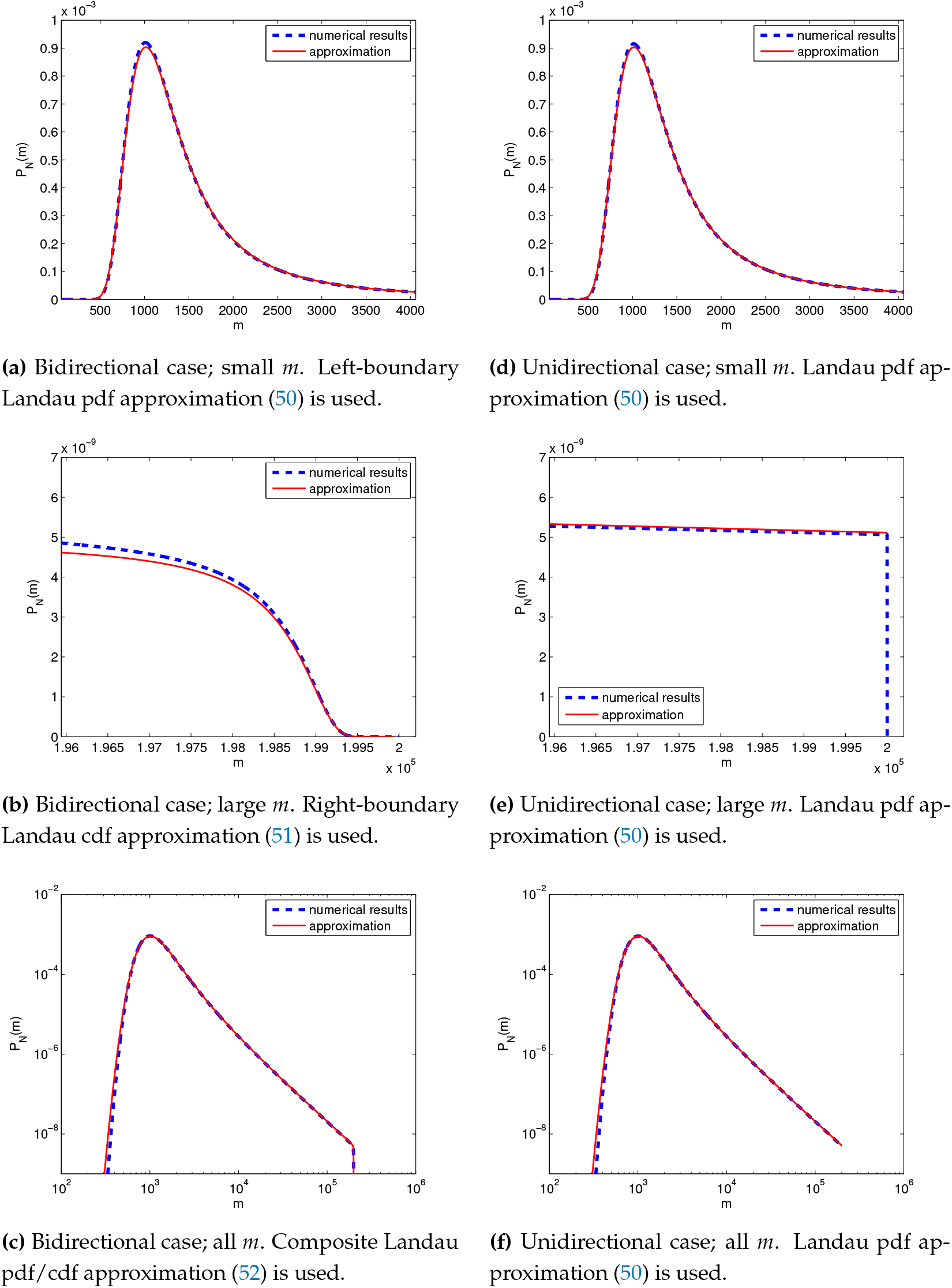
Probability *P*_*N*_ (*m*) of having *m* resistant cells in a total population of *N* cells as calculated numerically (blue dashed lines) and its asymptotic approximations (50), (51), (52) (red solid lines). The population starts from a single sensitive cell, and is left to grow until it consists of *N* = 2 *×* 10^5^ cells. **(a-c)** Bidirectional case: the probabilities of resistance acquisition and loss are *μ* = *λ* = 10^−3^. **(d-f)** Unidirectional case: *μ* = 10^−3^, *λ* = 0.

For a large number of resistant cells *m* in the population, we have an approximation (see Figure 3b)

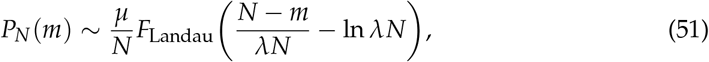

where 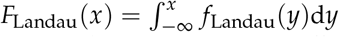 is the cumulative distribution function of the Landau distribution. The approximation follows from (41) for *N*_0_ = 1 and (47).

Finally, a composite approximation (Figure 3c)

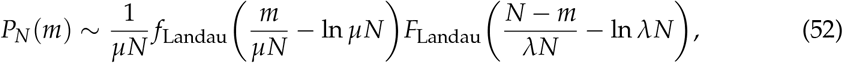

works well for all possible numbers *m* of sensitive cells. The approximation follows from (45) for *N*_0_ = 1 and (47).

In the irreversible, unidirectional, case (*λ* = 0), the Landau pdf (50) applies uniformly for the entire range 0 ≤*m* ≤ *N* of resistant subpopulation sizes (Figures 3d–3f). The probability distribution then exhibits a sharp cut-off at *m* = *N* (Figure 3e); contrastingly, the presence of a reverse transition smooths the cut-off into a Landau cdf in the right boundary layer (Figure 3b).

### 4.2. One sensitive and one resistant cell at the beginning

Mixed initial population develops a dramatically different dynamics than the homogeneous initial population. In an initial phase, which lasts until the population size reaches the reciprocal of the small resistance acquisition parameter, the dynamics are driven by the competition between the independent progeny of the two parent cells. A classical result implies that this tug of war will lead to a uniform distribution of resistant/sensitive cells. In this paper this classical result is incorporated in the “regular coarse-grained” approximation (20) with *m*_0_ = 1, *N*_0_ = 2. After the initial phase, boundary layers appear at the left and right corners of the interval *m ∈* {0, 1, …, *N*} due to resistance acquisition and loss. This paper provides nontrivial approximations of the boundary behaviour by the Landau cumulative distribution function.

For a small to moderate number of resistant cells *m* in the population, we have an approximation (Figure 4a)

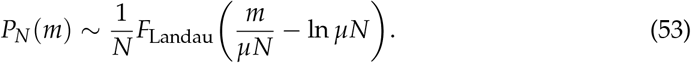

**Figure 4.**
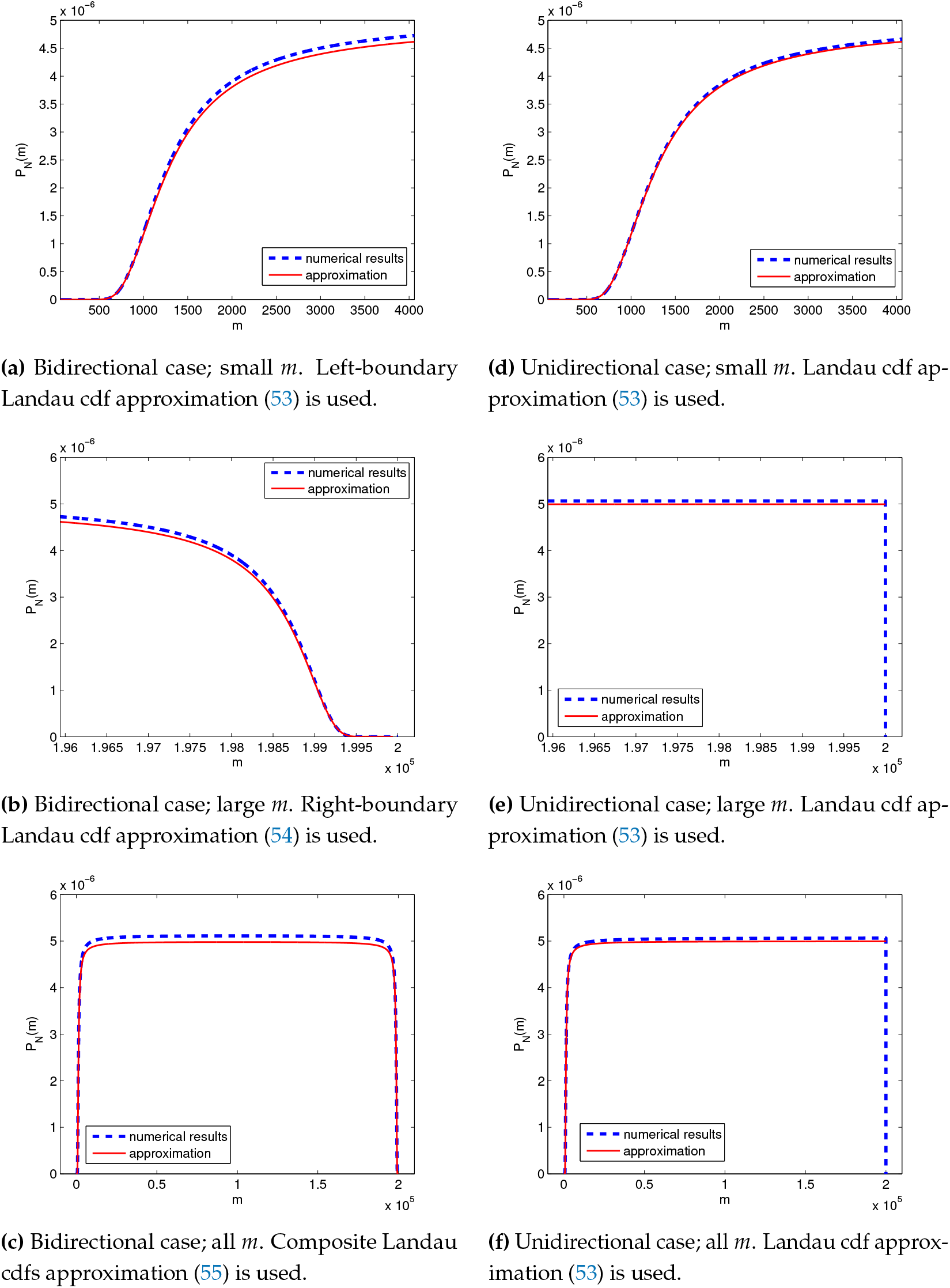
Probability *P*_*N*_ (*m*) of having *m* resistant cells in a total population of *N* cells as calculated numerically (blue dashed lines) and its asymptotic approximations (53), (54) and (55) (red solid lines). The population starts from two cells, one sensitive and one resistant, and is left to grow until it consists of *N* = 2 *×* 10^5^ cells. **(a-c)** Bidirectional case: The probabilities of resistance acquisition and loss are *μ* = *λ* = 10^−3^. **(d-f)** Unidirectional case: *μ* = 10^−3^, *λ* = 0.

The approximation is derived from (32) for *m*_0_ = 1, *N*_0_ = 2 and (47).

For a large number of resistant cells *m* in the population, we have an approximation (Figure 4b)

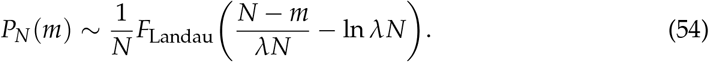

The approximation is derived from (40) for *m*_0_ = 1, *N*_0_ = 2 and (47).

Finally, just like in case of a sensitive initial population, also in case of a mixed initial population, one can find a composite approximation that works very well for all possible *m*. In this case it is given by (Figure 4c)

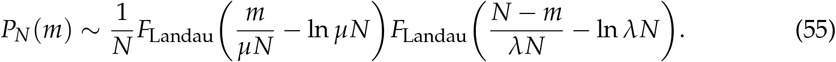

The approximation is derived from (44) for *m*_0_ = 1, *N*_0_ = 2 and (47).

In the unidirectional case (*λ* = 0), the Landau cdf approximation (50) applies uniformly for all resistant subpopulation sizes 0≤ *m* ≤ *N* (Figures 4d–4f). Again, a sharp cut-off is observed at *m* = *N* in the unidirectional case (Figure 4e), whereas a Landau cdf in the right boundary layer appears in the reversible case (Figure 4b).

### 4.3. More than one sensitive or resistant cell at the beginning

The cumulative distribution function is a partial summation/integration (up to a given level) of the probability mass/density function. The second partial summation/integration, third, etc., is defined recursively. The higher order summations/integrations of the Lea– Coulson/Landau density appear in the boundary behaviour of the process if the initial condition contains more than one cell of either type (resistant/sensitive). General formulae are provided in Sections 3.

### 4.4. Stochastic initial conditions

Our analysis so far concerned a deterministic initial condition (3). A previous study of a bidirectional model (based on different assumptions) [8] considered a stochastic initial condition, in which the population is derived from a single initial cell, which can be with a certain probability *ρ* sensitive and with the complementary probability it is resistant. By the superposition principle, the solution that is subject to the stochastic initial condition is equal to the weighted sum of the solution starting deterministically with one sensitive parent and the solution starting deterministically with one resistant parent. The Landau cumulative distribution functions in the tails of the two solutions are then negligible. For low to moderate values *m*, the probability of seeing *m* resistant cells among *N* total cells is approximated by

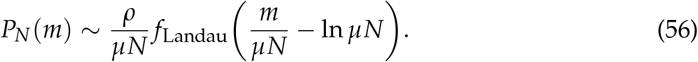

For a large number of resistant cells *m* in the population, we have an approximation

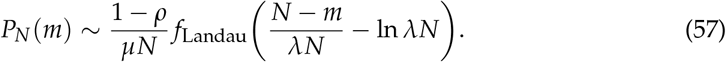

A uniformly valid solution is obtained by adding up (56) and (57):

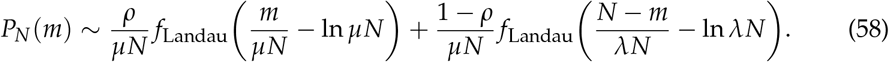

We note that this is a composite rather than log-composite approximation.

A special type of initial condition is if the switching between resistance and sensitivity is in balance: *ρ* = *λ*/(*μ* + *λ*), 1 − *ρ* = *μ*/(*μ* + *λ*). Then the mean resistant fraction satisfies 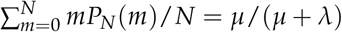, i.e. it does not change as the population grows [8].

### 4.5. Limitations of the approximations

The left boundary solution has the form of the Landau pdf/cdf located at *m* = *μN* ln *μN*, see Equations (50), (53), (56). For self-consistency, we must require that the boundary layer location be much less than the size of the entire definition domain, meaning that *μN* ln *μN*≪ *N*. This is seen to be equivalent to requiring ln *N*≪ 1/*μ*, where *μ*≪ 1. Thus, *N* should large, but not exponentially large, for the asymptotic approximations to apply.

## 5. Discussion

The paper focuses on a stochastic Luria–Delbrück fluctuation test model with reversible switching between two cellular states (for example, single cells being drug resistant or drug sensitive). We considered a fixed population ensemble formulation. Let us briefly discuss the corresponding fixed time ensemble version. In that, each cell has an exponentially distributed cell cycle duration with the same mean generation time regardless of the cell’s state. At the end of the cell cycle, the cell divides and produces a daughter cell, the state of which is chosen probabilistically as in the fixed ensemble formulation (Figure 1). The total population grows exponentially as per the Yule growth process with growth rate reciprocal to the mean generation time.

If one conditions on the Yule process reaching a given value *N*, one obtains the fixed ensemble model for the number of resistant cells as studied in this paper. Conversely, drawing *N* from the state of a Yule growth process at a given time *t*, one turns the fixed population ensemble to a fixed time ensemble formulation. Conditioning on a fixed population thus eliminates the proliferation noise. Another approach to this consists in calculating the relative fractions of sensitive/resistant cells [8].

Our model assumes that switching between sensitivity and resistance occurs at cell division events. The previous study [8] of a reversible switching case was based on different assumptions, which warrant a brief discussion. Like in the fixed time ensemble version of our model, the cell cycle is assumed to be exponentially distributed and the mean generation time is the same for both cell types. However, switching is uncoupled from the cell cycle in [8]: a daughter cell is always of same type as the mother cell, and any cell has a constant propensity to transition, at any time, to the opposite state.

Importantly, unlike in our model, there can be more than one switching event per cell cycle in the model of [8]. It is therefore expected, although this needs to be supported by further analysis, that the difference between the two formulations will be pronounced in the fast switching regime in which multiple switching events per cell cycle are probable. We note that in the model of [8], proliferation events increase the population by one, but the switching events do not; this makes the relationship between the fixed time and fixed population ensemble formulations more subtle.

The reversible model studied in this paper is a direct extension of the irreversible switching model as studied in [10]; indeed, setting to zero the probability *λ* that a resistant mother bears a sensitive daughter cell, one recovers the earlier model. The earlier study provides methods that are specifically applicable in the important regime of a large population subject to low-rate switching. We extended these methods to the current reversible case. We emphasise that the generalisation required nontrivial modifications in the methodology, notably in the systematic use of asymptotic matching between the small and large population solutions.

We note that the new approximation differs from that given in [10] only in a low probability distribution tail, as evidenced by Figure 3. Nevertheless, we believe that the intriguing appearance of the Landau cumulative distribution function in the tail will be interesting to a wide audience. Additionally, we have established that it also appears for mixed initial conditions, as illustrated in Figure 4.

## Author Contributions

Conceptualization, A.S.; methodology, P.B.; analysis P.B and A.H.; visualization, A.H.; writing—review and editing, P.B., A.H., and A.S. All authors have read and agreed to the published version of the manuscript.

## Institutional Review Board Statement

Not applicable

## Data Availability Statement

Not applicable

## Abbreviations

The following abbreviations are used in this manuscript:

Pmf: probability mass function
Pdf: probability density function
Cdf: cumulative distribution function
L–C: Lea–Coulson

## Appendix A.

### Master equation derivation

Our aim is to derive the relevant recurrent equation for *P*_*N*+1_(*m*), if the probability *P*_*N*_(*m*) for a population of size *N* and for all possible *m* is known. In other words, we want to express the probability that after one birth event, the size of the resistant subpopulation would stay *m*. One may realize that the sought probability *P*_*N*+1_(*m*) can be expressed as a sum of probabilities *A*_*i*_, *i* = 1, 2, 3, 4 corresponding to four disjunctive situations that can happen and that are depicted on schematic Fig. 1:

1. *Parent cell sensitive, descendant resistant:* 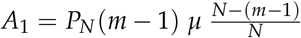
2. *Parent cell sensitive, descendant sensitive:* 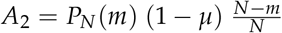
3. *Parent cell resistant, descendant resistant:* 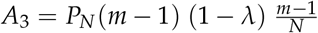
4. *Parent cell resistant, descendant sensitive:* 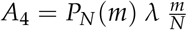

The sum *A*_1_ + *A*_2_ + *A*_3_ + *A*_4_ provides the right hand side of the master equation (1).

## Appendix B.

### Polya urn models

Suppose that we have an urn that consists of *w* white balls and *b* black balls at the beginning. Then we withdraw a ball from the urn one at a time, look at its color, return the ball to the urn and add *c* balls of the same color. The probability that *x* white balls are drawn in a sample of *n* withdrawals is

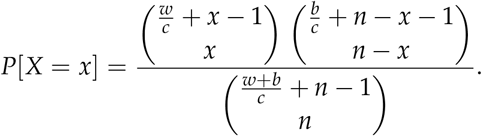

Also, the number of balls of a specific colour in the Polya urn model follows the negative hypergeometric distribution [24].

We can adapt Polya urn model to our situation in the following way: Firstly, we realize that *c* = 1 in our case, as in every birth event, only one new cell is born. Furthermore, let *w* and *b* stand for the number of resistant, respectively sensitive, cells in the initial population. If *n* denotes the number of birth events and *x* denotes the number of sensitive cells from *n* divided, than the following relations with previous notation hold

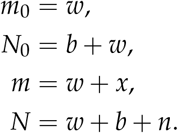

Therefore, we can write that as *μ* (and *λ*) tend to zero,

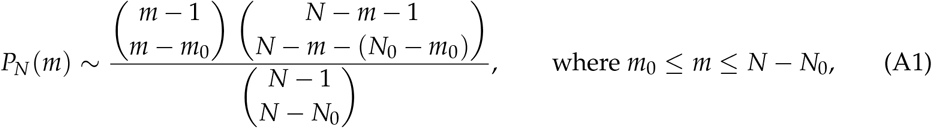

where the tilde sign is understood in the usual sense of asymptotic expansions. Rearrangement of combinatorial numbers yields (4).

## Appendix C.

### Computations

Symbolic computation of the sum (9) using software Matlab.

~~~
syms L N m N_0
A = factorial(N-L -1);
B = factorial(N-L-m);
s = symsum (L* A/B, L, N_0, N-m);
assume (m >0);
simplify (s)
~~~

## References

1. Luria, S.E.; Delbrück, M. Mutations of bacteria from virus sensitivity to virus resistance. Genetics 1943, 28, 491.

2. Shaffer, S.M.; Dunagin, M.C.; Torborg, S.R.; Torre, E.A.; Emert, B.; Krepler, C.; Beqiri, M.; Sproesser, K.; Brafford, P.A.; Xiao, M.; et al. Rare cell variability and drug-induced reprogramming as a mode of cancer drug resistance. Nature 2017, 546, 431–435.

3. Shaffer, S.M.; Emert, B.L.; Hueros, R.A.R.; Cote, C.; Harmange, G.; Schaff, D.L.; Sizemore, A.E.; Gupte, R.; Torre, E.; Singh, A.; et al. Memory sequencing reveals heritable single-cell gene expression programs associated with distinct cellular behaviors. Cell 2020, 182, 947–959.

4. Hossain, T.; Singh, A.; Butzin, N.C. Escherichia coli cells are primed for survival before lethal antibiotic stress. bioRxiv 2022.

5. Chang, C.A.; Jen, J.; Jiang, S.; Sayad, A.; Mer, A.S.; Brown, K.R.; Nixon, A.M.; Dhabaria, A.; Tang, K.H.; Venet, D.; et al. Ontogeny and vulnerabilities of drug-tolerant persisters in her2+ breast cancer. Cancer discovery 2022, 12, 1022–1045.

6. Harmange, G.; Hueros, R.A.R.; Schaff, D.L.; Emert, B.L.; Saint-Antoine, M.M.; Nellore, S.; Fane, M.E.; Alicea, G.M.; Weeraratna, A.T.; Singh, A.; et al. Disrupting cellular memory to overcome drug resistance. bioRxiv 2022.

7. Saint-Antoine, M.M.; Grima, R.; Singh, A. A fluctuation-based approach to infer kinetics and topology of cell-state switching. bioRxiv 2022.

8. Saint-Antoine, M.M.; Singh, A. Moment-Based Estimation of State-Switching Rates in Cell Populations. bioRxiv 2022.

9. Zheng, Q. Progress of a half century in the study of the Luria–Delbrück distribution. Mathematical biosciences 1999, 162, 1–32.

10. Kessler, D.A.; Levine, H. Large population solution of the stochastic Luria–Delbrück evolution model. Proceedings of the National Academy of Sciences 2013, 110, 11682–11687.

11. Keller, P.; Antal, T. Mutant number distribution in an exponentially growing population. Journal of Statistical Mechanics: Theory and Experiment 2015, 2015, P01011.

12. Nicholson, M.D.; Antal, T. Universal asymptotic clone size distribution for general population growth. Bulletin of Mathematical Biology 2016, 78, 2243–2276.

13. Antal, T.; Krapivsky, P. Exact solution of a two-type branching process: models of tumor progression. Journal of Statistical Mechanics: Theory and Experiment 2011, 2011, P08018.

14. Angerer, W.P. An explicit representation of the Luria–Delbrück distribution. Journal of mathematical biology 2001, 42, 145–174.

15. Kessler, D.A.; Levine, H. Scaling solution in the large population limit of the general asymmetric stochastic Luria–Delbrück evolution process. Journal of statistical physics 2015, 158, 783–805.

16. Jolly, C.; Cook, A.; Raferty, J.; Jones, M. Measuring bidirectional mutation. Journal of theoretical biology 2007, 246, 269–277.

17. Sorace, R.; Komarova, N.L. Accumulation of neutral mutations in growing cell colonies with competition. Journal of theoretical biology 2012, 314, 84–94.

18. Cheek, D.; Antal, T. Genetic composition of an exponentially growing cell population. Stochastic Processes and their Applications 2020, 130, 6580–6624.

19. Lea, D.E.; Coulson, C.A. The distribution of the numbers of mutants in bacterial populations. Journal of genetics 1949, 49, 264–285.

20. Pakes, A.G. Remarks on the Luria–Delbrück distribution. Journal of Applied Probability 1993, 30, 991–994.

21. Angerer, W.P. A note on the evaluation of fluctuation experiments. Mutation Research/Fundamental and Molecular Mechanisms of Mutagenesis 2001, 479, 207–224.

22. Nolan, J.P. Univariate stable distributions; Springer, 2020.

23. Murray, J.D. Mathematical biology: I. An introduction; Springer, 2002.

24. Johnson, N.L.; Kotz, S.; Kemp, A.W. Univariate discrete distributions; John Wiley & Sons, 2005.

25. Kevorkian, J.; Cole, J.D. Perturbation methods in applied mathematics; Springer Science & Business Media, 2013.

26. Mahmoud, H. Pólya urn models; Chapman and Hall/CRC, 2008.

27. Walczak, A.M.; Mugler, A.; Wiggins, C.H. Analytic methods for modeling stochastic regulatory networks. Computational Modeling of Signaling Networks 2012, pp. 273–322.

28. Landau, L.D. On the energy loss of fast particles by ionization. J. Phys. 1944, 8, 201–205.

